# Widespread lateral gene transfer among grasses

**DOI:** 10.1101/2020.02.17.952150

**Authors:** Samuel G. S. Hibdige, Pauline Raimondeau, Pascal-Antoine Christin, Luke T. Dunning

## Abstract

- Lateral gene transfer (LGT) has been documented in a broad range of prokaryotes and eukaryotes, and it can promote adaptation. LGT of functional nuclear genes has been reported among some plants, but systematic studies are needed to assess the frequency and facilitators of LGT in the group.
- We scan the genomes of a diverse set of 17 grass species that span more than 50 million years of divergence and include major crops to identify grass-to-grass protein-coding LGT.
- We identify LGT in 13 species, with significant variation in the amount each received. Rhizomatous species acquired statistically more genes, probably because this growth habit boosts opportunities for transfer into the germline. In addition, the amount of LGT increases with phylogenetic relatedness, which might reflect genomic compatibility amongst close relatives facilitating successful transfers. However, genetic exchanges among highly divergent species with overlapping distributions also occur, pointing to an additional role of biogeography.
- Overall, we show that LGT is a widespread phenomenon in grasses, which has moved functional genes across the entire grass family into domesticated and wild species alike. The dynamics of successful LGT appears to be dependent on both opportunity (co-occurrence and rhizomes) and compatibility (phylogenetic distance).

## 1. Introduction

The adaptive potential of a species is limited by its evolutionary history, the amount of standing genetic variation and the rate of new mutations (Barrett & Schluter, 2008). Lateral gene transfer (LGT) enables organisms to overcome these limitations by exchanging genetic material between lineages that have evolved significant reproductive barriers (Doolittle, 1999). LGT is an important evolutionary force in prokaryotes, with up to 60% of genes within a species pan-genome being acquired in this manner (Freschi *et al*., 2018). The genes transferred can have a dramatic effect on adaptation, facilitating the colonisation of new niches and the development of novel phenotypes, as exemplified by the rapid spread of antibiotic resistance in bacteria (Ochman *et al*., 2000). While LGT is more prevalent in prokaryotes, it has also been documented in a variety of multicellular eukaryotes (reviewed in: Anderson, 2005; Keeling & Palmer, 2008; Schönknecht *et al*., 2014; Husnik *et al*., 2018; Van Etten & Bhattacharya, *in press*), including plants (reviewed in: Richardson & Palmer, 2007; Gao *et al*., 2014; Wickell & Li, 2019).

DNA has been transferred into plants from prokaryotes, fungi and viruses, with recipients in particular in algae (Cheng *et al*., 2019) and bryophytes (Yue *et al*., 2012; Maumus *et al*., 2014; Bowman *et al*., 2017; Zhang *et al*., 2020). Concerning plant-to-plant transfers, a majority of nuclear LGT reported so far involve the transfer of genetic material between parasitic species and their hosts, with examples from the genera *Cuscuta* (Vogel *et al*., 2018; Yang *et al*., 2019), *Rafflesia* (Xi et al., 2012), and *Striga* (Yoshida et al., 2010). However, plant-to-plant LGT is not restricted to parasitic interactions, and it has been recorded in ferns (Li *et al*., 2014) and eight different species of grass (Vallenback *et al*., 2008; Christin *et al*., 2012a; Prentice *et al*., 2015; Mahelka *et al*., 2017; Dunning *et al*., 2019). Grasses represent one of the best systems to investigate factors promoting LGT between non-parasitic plants as they are the only group where multiple LGT recipients have been identified, and there is extensive genomic resources available due to their economic and ecological importance (Chen *et al*., 2018). Early examples of grass-to-grass LGT were largely obtained incidentally, and only one grass genome (*Alloteropsis semialata*) has been comprehensively scanned, with 59 LGTs identified using stringent phylogenetic filters (Dunning *et al*., 2019). These 59 protein-coding genes were transferred from at least nine different donors as part of 23 large fragments of foreign DNA (up to 170 kb). A majority of the acquired LGTs within *A. semialata* are expressed, with functions associated with photosynthesis, disease resistance and abiotic stress tolerance (Dunning *et al*., 2019; Phansopa *et al., In press*). While reports of LGT in other species in the group suggest it is a widespread phenomenon, its full distribution within the family remains to be assessed.

Grasses are very diverse (Soreng *et al*., 2015), with more than 12,000 species exhibiting extensive phenotypic variation that may influence LGT dynamics. In particular, the family contains both annuals and perennials. If LGT happens during vegetative growth (e.g. root-to-root inosculation), the number of LGT is predicted to be higher in perennial and rhizomatous species. Conversely, if LGT happens through illegitimate pollination, the number of LGT may not vary with growth form or be higher in annuals that produce seeds more frequently. Finally, successful transfers might be more likely to occur between closely-related groups with similar genome features as observed in prokaryotes (Skippington & Ragan, 2012; Soucy *et al*., 2015). Most of the grass diversity is clustered in the two BOP and PACMAD sister groups that diverged more than 50 million years ago (Christin *et al*., 2014). Each of the two groups has more than 5,000 taxa and includes model species with complete genomes (Soreng *et al*., 2015). The family therefore offers unparalleled opportunities to determine whether functional characteristics or phylogenetic distance determine the amount of LGT among non-parasitic plants.

In this study, we use a phylogenomic approach to scan 17 different grass genomes and quantify LGT among them. The sampled species belong to five different clades of grasses, two from the BOP clade (Oryzoideae and Pooideae) and three from the PACMAD clade (Andropogoneae, Chloridoideae, and Paniceae). Together, these five groups contain more than 8,000 species or over 70% of the diversity within the whole family (Soreng *et al*., 2015). In our sampling, each of these five groups is represented by at least two divergent species, allowing us to monitor the number of transfers among each group. In addition, the species represent a variety of domestication statuses, life-history strategies, genome sizes, and ploidy levels (Table 1). Using this sampling design, we (i) test whether LGT is more common in certain phylogenetic lineages, and (ii) test whether some plant characters are associated with a statistical increase of LGT. We then focus on the donors of the LGT received by the Paniceae tribe, a group for which seven genomes are available, to (iii) test whether the number of LGT increases with phylogenetic relatedness. Our work represents the first systematic quantification of LGT among members of a large group of plants and sheds new light on the conditions that promote genetic exchanges across species boundaries in plants.

**Table 1:** Model species used in this study and associated traits.

| Clade | Species | Ploid. | 1C | #Genes tested | Cult. | LH | Clim. | Phot. | Cont. | Rhiz. |
| --- | --- | --- | --- | --- | --- | --- | --- | --- | --- | --- |
| Pooideae | *Brachypodium distachyon*<sup>1</sup> | 2n | 0.31 | 17204 | N | A | Temp | C<sub>3</sub> | 6 | N |
| Pooideae | *Hordeum vulgare*<sup>2</sup> | 2n | 5.39 | 16192 | Y | A | Temp | C<sub>3</sub> | 6 | N |
| Pooideae | *Triticum aestivum*<sup>3</sup> | 6n | 16.95 | 56619 | Y | A | Temp | C<sub>3</sub> | 6 | N |
| Oryzoideae | *Oryza sativa*<sup>4</sup> | 2n | 0.49 | 19259 | Y | A | Trop | C<sub>3</sub> | 6 | N |
| Oryzoideae | *Leersia perrieri*<sup>5</sup> | 2n | 0.32 | 15777 | N | A | Trop | C<sub>3</sub> | 1 | N |
| Chloridoideae | *Eragrostis tef*<sup>6</sup> | 4n | 0.69 | 30605 | Y | A | Temp | C<sub>4</sub> | 5 | N |
| Chloridoideae | *Oropetium thomaeum*<sup>7</sup> | 2n | 0.29 | 15168 | N | A | Temp | C<sub>4</sub> | 2 | N |
| Chloridoideae | *Zoysia japonica*<sup>8</sup> | 4n | 0.42 | 20416 | Y | P | Temp | C<sub>4</sub> | 1 | Y |
| Andropogoneae | *Sorghum bicolor*<sup>9</sup> | 2n | 0.69 | 21962 | Y | A | Temp | C<sub>4</sub> | 6 | N |
| Andropogoneae | *Zea mays*<sup>10</sup> | 2n | 2.65 | 25866 | Y | A | Temp | C<sub>4</sub> | 6 | N |
| Paniceae | *Alloteropsis semialata*<sup>11</sup> | 2n | 1.10 | 23071 | N | P | Trop | C<sub>4</sub> | 3 | Y |
| Paniceae | *Cenchrus americanus*<sup>12</sup> | 2n | 2.65 | 20159 | Y | A | Temp | C<sub>4</sub> | 4 | N |
| Paniceae | *Dichanthelium oligosanthos*<sup>13</sup> | 2n | 0.96 | 17761 | N | P | Cold | C<sub>3</sub> | 1 | N |
| Paniceae | *Echinochloa crus-galli*<sup>14</sup> | 6n | 1.37 | 54181 | N | A | Temp | C<sub>4</sub> | 6 | N |
| Paniceae | *Panicum hallii*<sup>15</sup> | 2n | 0.55 | 30255 | N | P | Temp | C<sub>4</sub> | 1 | N |
| Paniceae | *Panicum virgatum*<sup>16</sup> | 4n | 1.89 | 45043 | Y | P | Cold | C<sub>4</sub> | 3 | Y |
| Paniceae | *Setaria italica*<sup>17</sup> | 2n | 0.49 | 27465 | Y | A | Temp | C<sub>4</sub> | 6 | N |

Ploid. = Ploidy; 1C = 1C genome size in Gb; Cult. = cultivated (Y = yes; N = no); LH = life history (A = annual; P = perennial); Clim. = climate (Temp = temperate; Trop = tropical); Phot. = photosynthetic type; Cont. = number of continents; Rhiz. = rhizomatous (Y = yes; N = no).
<sup>1</sup>International Brachypodium Initiative, 2010; <sup>2</sup>International Barley Genome Sequencing Consortium, 2012; <sup>3</sup>International Wheat Genome Sequencing Consortium, 2014; <sup>4</sup>Goff *et al.*, 2002; <sup>5</sup>Stein *et al.*, 2018; <sup>6</sup>Cannarozzi *et al.*, 2014; <sup>7</sup>VanBuren *et al.*, 2015; <sup>8</sup>Tanaka *et al.*, 2016; <sup>9</sup>Patterson *et al.*, 2009; <sup>10</sup>Schnable *et al.*, 2009; <sup>11</sup>Dunning *et al.*, 2019; <sup>12</sup>Varshney *et al.*, 2017; <sup>13</sup>Studer *et al.*, 2016; <sup>14</sup>Guo *et al.*, 2017; <sup>15</sup>Lovell *et al.*, 2018; <sup>16</sup>*Panicum virgatum* v4.1, DOE-JGI, http://phytozome.jgi.doe.gov/; <sup>17</sup>Bennetzen *et al.*, 2012

## 2. Materials and Methods

### 2.1 Detecting grass-to-grass LGT

We modified the approach previously used by Dunning *et al*., (2019) to identify grass-to-grass LGT. Specifically, we did not use the initial mapping filtering step from Dunning *et al*. as it relied on the availability of high-coverage genome data for pairs of closely related species. In total, 17 genomes were scanned for LGT (Table 1), with all phylogenetic analyses based on coding sequences (total = 817,621 genes; mean per species 48,095 genes; SD = 26,764 genes).

In the first step, 37-taxa trees were constructed using data from the 17 grass genomes (Table 1), supplemented with transcriptome data for 20 additional species from across the grass family (Moreno-Villena *et al*., 2018; Table S1). BLASTn was used to identify the top-match for each gene in the 36 other species with a minimum match length of 300bp (not necessarily a single continuous BLAST match). Nucleotide alignments were generated by aligning the BLASTn matching regions to the query sequence using the ‘add fragments’ parameter in MAAFT v7.427 (Katoh & Standley, 2013). If the BLASTn match for a species was fragmented, the different fragments were joined into a single sequence after they had been aligned. Alignments with less than ten species were considered non informative and consequently discarded (retained 55.9% of genes; total = 457,003 genes; mean per species 26,883 genes; SD = 13,042 genes; Table S2). For each alignment with ten species or more, a maximum-likelihood phylogenetic tree was inferred using PhyML v.20120412 (Guindon & Gascuel, 2003) with the GTR+G+I substitution model. Each topology was then midpoint rooted using the phytools package in R and perl scripts were used to identify genes from each focus species nested within a different group of grasses. We focused on five groups (Andropogoneae, Chloridoideae, Oryzoideae, Paniceae and Pooideae) represented by at least two complete genomes that were supported by most gene trees in a previous mulitgene coalescent species tree analysis (Figure 1; Dunning *et al*., 2019). The whole set of analyses were later repeated to detect LGT between well supported subclades within the Paniceae, considering LGT received from two clades represented by two genomes and supported by most gene trees in previous analyses (i.e. Cenchrinae and Panicinae, Figure 1). To be considered as nested, the sister species of the query gene, and their combined sister group, had to belong to the same grass group to which the query gene does not belong. For genes that were nested, the analysis was repeated with 100 bootstrap replicates to verify that the nesting of the query sequence was supported by a minimum of 50% of the bootstrap replicates.

**Fig 1:**
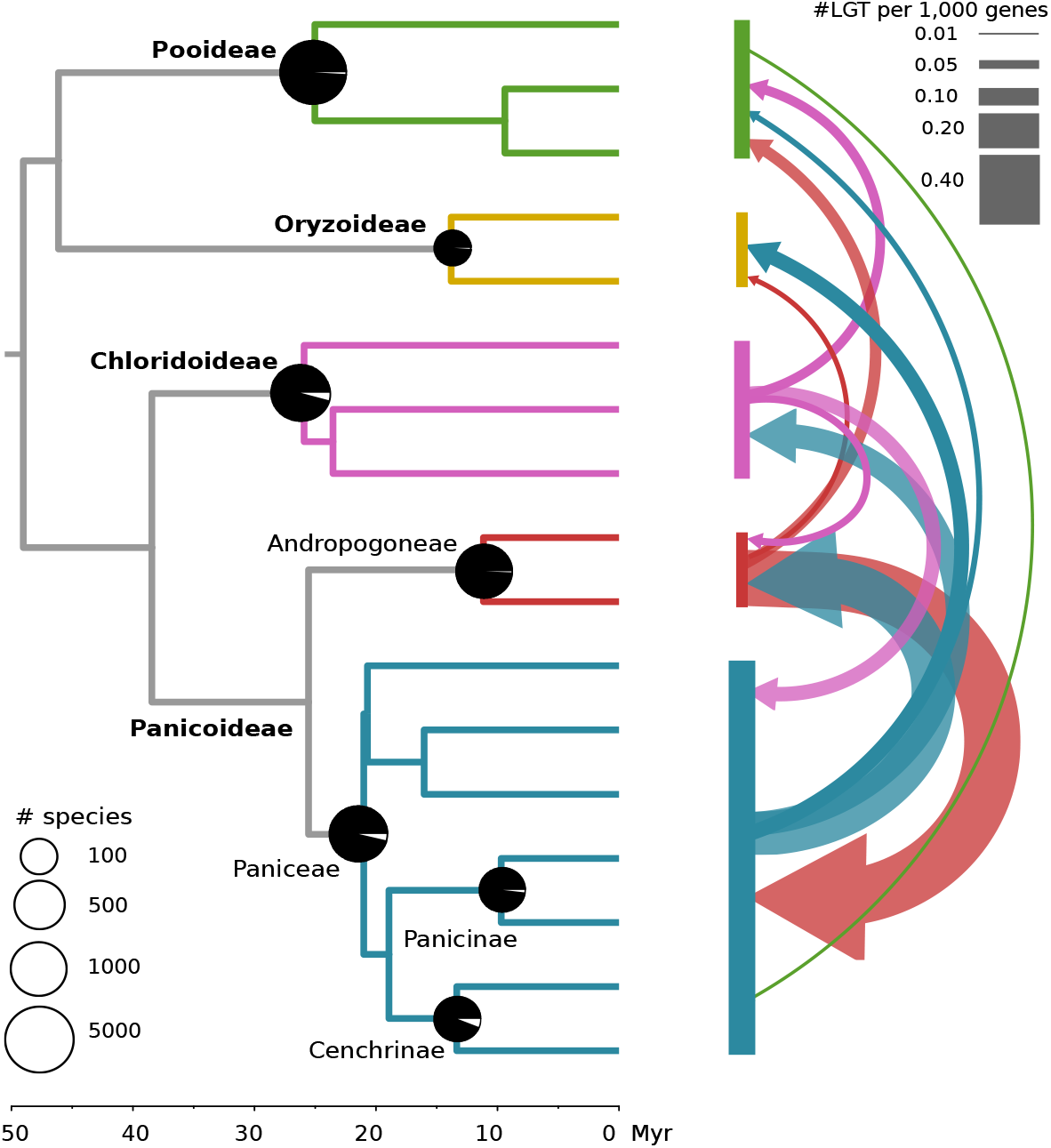
Distribution of lateral gene transfers among grasses. Time-calibrated phylogenetic tree of 17 model grass species used in this study (extracted from Christin et al., 2014; scale in million years - Myr). The direction of LGT between grass clades is shown with arrows whose size is proportional to the number of LGT received. The black portion of pie charts on key nodes of the phylogeny indicates the quartet support for the observed topology based on a multigene coalescence analysis (Dunning et al., 2019). The size of each pie chart is proportional to the number of species within the clade (Soreng et al., 2015). Numbers at the tips are the number of LGT detected in each genome.

The second filtering step was applied to those candidates that passed the first phylogenetic filter. For those, we performed a second round of filtering using data from 105 genome/transcriptome datasets belonging to 85 species, including the datasets used for the 37-taxa trees (Table S1). For each LGT candidate, we used BLASTn to identify all matches (not just the best match) with a minimum alignment length of 300bp (not necessarily a single continuous blast match) in each of the 105 datasets. Alignments were generated as previously, before being re-aligned as codons using MAAFT and manually trimmed with a codon-preserving method to remove poorly aligned regions. Maximum likelihood phylogenies were then inferred using PhyML v.21031022, with the best substitution model identified by Smart Model Selection SMS v.1.8.1 (Lefort *et al*., 2017). The trees were manually inspected and discarded if: i) there were less than three species within or outside the LGT donor clade; ii) the LGT candidate was not nested within another group of grasses with the increased taxon sampling; or iii) the tree had obvious paralogy problems due to gene duplication events. For retained candidates, we removed paralogs representing duplicates originating before the core grasses (BOP and PACMAD clades; Soreng *et al*., 2015), and joined fragmented transcripts from a single data set if they were nested within the same phylogenetic group. The tree inference was then repeated with 100 bootstraps, and the trees were again manually inspected, discarding candidates where the nesting was supported by <70% bootstrap replicates. Finally, BLASTx was used to annotate the LGT candidates against the SwissProt database.

After these two successive filters, retained candidates were subjected to further validation. To verify the nesting of the candidate LGT was not due to convergent adaptive amino acid substitutions, we generated phylogenetic trees based solely on 3^rd^ codon positions, which are less subject to positive selection (Christin *et al*., 2012b). Phylogenetic trees were generated as above and were manually inspected to confirm the LGT scenario. To verify that the LGT scenario was statistically better than the species tree, we then conducted approximately unbiased (AU) topology tests that compared the maximum likelihood topology with a topology representing the null hypothesis (forcing monophyly of the donor and recipient clades; recipients for the within-Paniceae analysis were constrained at the genus level if they did not belong to the Cenchrinae or Panicinae). The null topology was inferred by first constraining the clades and inferring a tree with the GTR + G model in RaxML v.8.2.12 (Stamatakis, 2014), before using this topology as a constraint for a maximum likelihood phylogeny inferred with PhyML as described above. The AU tests were then performed in Consel v.1.20 (Shimodaira & Hasegawa, 2001) using the site-wise likelihood values generated by PhyML, and p-values were Bonferroni corrected to account for multiple testing. LGTs with non-significant results (p-value > 0.05) were discarded. In some cases, no native copy was present in any species from the group containing the focus species, preventing AU tests. These genes were retained, although the numbers were recorded separately (Table 2; n.b. statistics reported and values quoted in the text include these genes).

**Table 2:** Number of lateral gene transfers (LGT) detected between the five clades.

| Clade | Species | # LGT | Donor clade |  |  |  |  |
| --- | --- | --- | --- | --- | --- | --- | --- |
|  |  |  | Pooid. | Ory. | Chlor. | Andro. | Pan. |
| Pooideae | *Brachypodium distachyon* | 4 | - | 0 | 0 | 4 | 0 |
| Pooideae | *Hordeum vulgare* | 0 | - | 0 | 0 | 0 | 0 |
| Pooideae | *Triticum aestivum* | 8(10) | - | 0 | 5 | 0(2) | 3 |
| Oryzeae | *Oryza sativa* | 0 | 0 | - | 0 | 0 | 0 |
| Oryzeae | *Leersia perrieri* | 1(4) | 0 | - | 0 | 1 | 0(3) |
| Chloridoideae | *Eragrostis tef* | 1(9) | 0 | 0 | - | 0 | 1(9) |
| Chloridoideae | *Oropetium thomaeum* | 0 | 0 | 0 | - | 0 | 0 |
| Chloridoideae | *Zoysia japonica* | 0 | 0 | 0 | - | 0 | 0 |
| Andropogoneae | *Sorghum bicolor* | 2(3) | 0 | 0 | 0 | - | 2(3) |
| Andropogoneae | *Zea mays* | 11 | 0 | 0 | 2 | - | 9 |
| Paniceae | *Alloteropsis semialata* | 20 | 0 | 0 | 4 | 16 | - |
| Paniceae | *Cenchrus americanus* | 15 | 0 | 0 | 5 | 10 | - |
| Paniceae | *Dichanthelium oligosanthos* | 4 | 4 | 0 | 0 | 0 | - |
| Paniceae | *Echinochloa crus-galli* | 10 | 0 | 0 | 3 | 7 | - |
| Paniceae | *Panicum hallii* | 8 | 0 | 0 | 6 | 2 | - |
| Paniceae | *Panicum virgatum* | 30 | 0 | 0 | 1 | 29 | - |
| Paniceae | *Setaria italica* | 7 | 0 | 0 | 0 | 7 | - |

The numbers in parentheses include genes for which approximate unbiased (AU) topology tests could not be performed as no native copy from the same clade was present to constrain the tree topology. Pooid. = Pooideae; Ory. = Oryzoideae; Chlor. = Chloridoideae; Andro. = Andropogoneae; Pan. = Paniceae.

For candidates retained after these extra validation steps, new phylogenetic trees were inferred with a denser species sampling to refine the identification of the potential donor. Illumina short-read data sets (n = 71; 65 sp.; Table S1) were added to the trees using the method described in Dunning *et al*., (2019). The dense trees were then manually inspected and any presenting strong discrepancies with the expected species relationships were discarded.

In summary, To be considered as an LGT each gene (i) had to be nested within one of the other four groups of grasses (Figure 1); (ii) their nesting had to be well supported (≥ 70% bootstrap support); (iii) potential parology problems had to be ruled out (i.e. discarding phylogenies with multiple apparent duplication events that can explain the phylogenetic incongruence); (iv) the nesting had to be supported by phylogenetic trees constructed solely from the 3rd codon positions, which are less subject to adaptive convergent evolution; and (v) where possible, the nesting had to be supported by approximately unbiased (AU) tests to confirm the LGT topology was a significantly better fit than a topology constrained to match the species tree (see Figure 2 for exemplar LGT). Phylogenetic trees and alignments are included as supplementary datasets. All analyses were preformed using publicly available data (Table S1).

**Fig 2:**
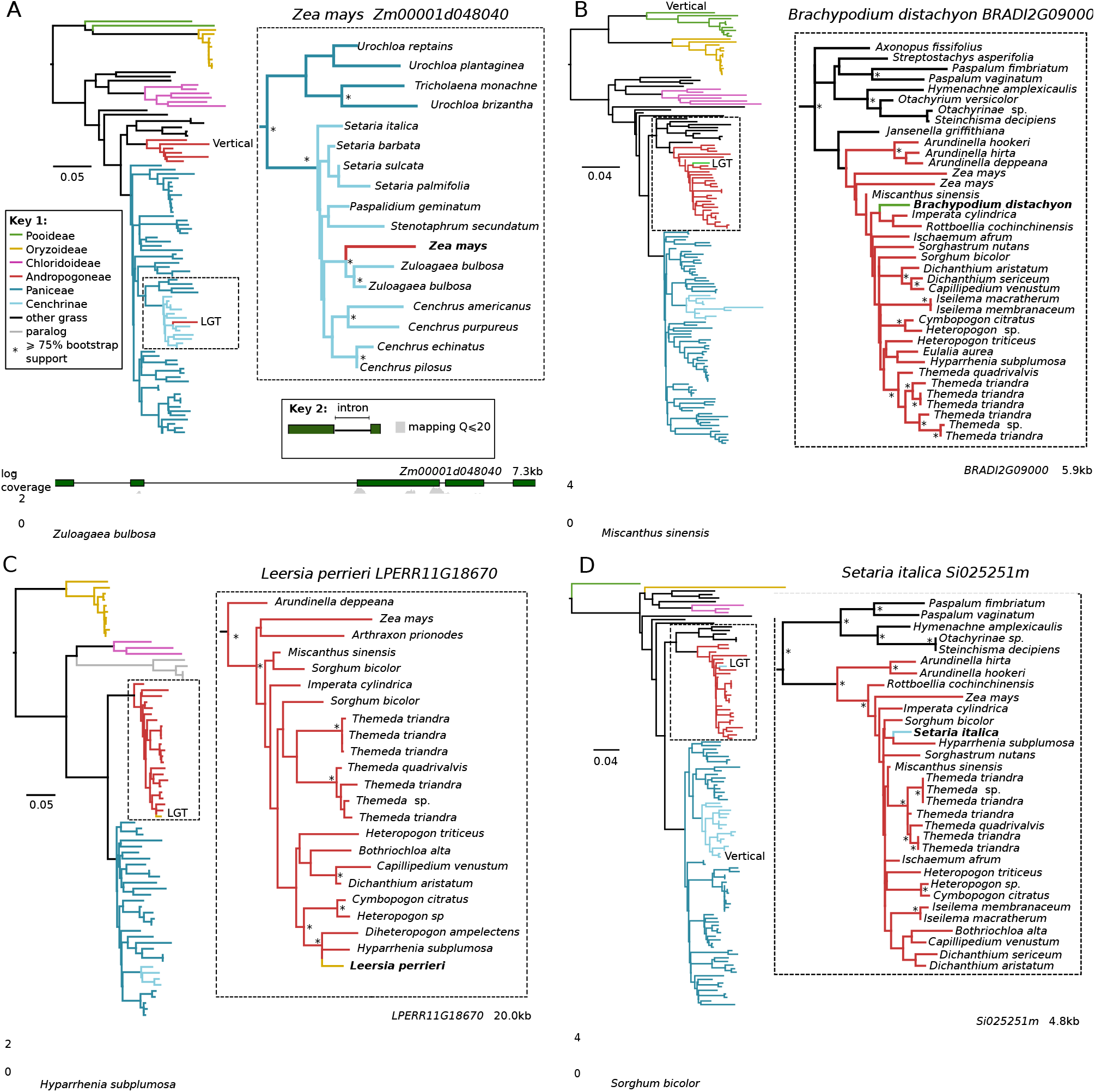
Four examples of grass-to-grass lateral gene transfer. Each panel (A-D) shows an exemplar grass-to-grass LGT, with full and expanded regions of maximum likelihood phylogenies shown. A coverage plot for each gene model is shown below, generated from short-read mapping data for a species closely related to the LGT donor.

### 2.2 Synteny analyses

Synteny analyses were performed for a subset of seven diploid grass genomes with high-quality reference genomes (*Brachypodium distachyon, Oryza sativa, Oropetium thomaeum, Panicum hallii, Setaria italica, Sorghum bicolor*, and *Zea mays*) using SynFind (Tang *et al*., 2015). For each LGT in these species, we determined whether genes from the other reference genomes identified as orthologs to the native copy in the phylogenetic trees were syntenic to the LGT or the native copy based on the highest syntelog score (Table S3).

### 2.3 Analyses of replicate sequencing runs to check for potential contamination

Independently sequenced runs for each of the model species were screened for the presence of each LGT, as potential contaminations would not appear in multiple replicates. Paired-end Illumina whole-genome data were obtained from NCBI Sequence Read Archive and mapped to the reference genome using bowtie2 v.2.3.5.1 (Langmean & Salzberg, 2012). Mean coverage depths for the coding sequence of each gene in the genome were then calculated using bedtools v2.26.0 (Quinlan & Hall, 2010), with large bam files down-sampled with Picard Tools v.2.13.2-SNAPSHOT (Broad Institute, 2019).

### 2.4 Grass traits

Plant traits were obtained from a variety of sources. Life history, distribution, growth form and the domestication status were retrieved from GrassBase (Clayton *et al*., 2016). 1C Genome sizes were obtained from the Plant DNA C-values database (Bennett & Leitch, 2005), and climatic information from Watcharamongkol *et al*., (2018). The climate data for *Oropetium thomaeum* was not included in Watcharamongkol *et al*., (2018), and was therefore retrieved from GBIF (GBIF.org; 11th July 2019) GBIF Occurrence Download https://doi.org/10.15468/dl.wyhtoo) and WorldClim (Harris *et al*., 2014; Fick & Hijmans, 2017) using the same methods. All statistical tests were preformed in R v.3.0.2, with the expected frequencies for chi-square tests based on the number of genes tested within each species (Table 1). The Kruskal-Wallis tests performed using absolute LGT numbers, which were divided into donor groups when testing whether some clades were more frequent donors than others. To determine if any trait or genome feature was associated with the number of LGT, we preformed phylogenetic generalized least squares (PGLS) to account for the relatedness between samples. The PGLS analysis was preformed in R with the ‘caper’ package (Orme *et al*., 2013) using a time-calibrated phylogenetic tree retrieved from Christin *et al*., (2014), and various traits as explanatory variables (Table 1). Individual and iterative models were performed, removing the least significant variable until only significant variables remained (p-value <0.05).

## 3. Results

### 3.1 LGT occurs in all lineages and functional types of grass

Out of the 817,612 grass genes from the 17 grass genomes (Table 1) screened, 55.89% had sufficient homologous grass sequences (≥ 10 taxa) for reliable phylogenetic reconstruction (Table 2 & Table S2), and were tested for LGT. A majority (99.73%) of the initial 37-taxa phylogenies did not support a scenario of LGT among the five grass groups, with successive filtering resulting in the identification of 135 LGT candidates across the 17 grass genomes in this initial analysis (Table 2; full results Table S2). The number of LGT received varied among species (p-value < 0.01; Chi-square test; mean = 8.4; SD=9.0; range=0 - 34; Table S2), with the highest numbers observed in *Panicum virgatum* (n= 30), *Alloteropsis semialata* (n=20), and *Cenchrus americanus* (n=15). It should be noted that only a subset of the 59 previously reported LGT in *Alloteropsis semialata* (Dunning *et al*., 2019) are retrieved as the previous analysis examined additional groups of donors not considered here, and secondary candidates based solely on read-mapping patterns were not recorded in the present study. Despite the significant variation between species, the difference among the five phylogenetic groups was not significant (p-value = 0.16, Kruskal-Wallis test). Overall, our results show that LGT is widespread across the grass family and occurs in a majority of species (Figure 1; Table 2). No LGT were detected in four of the 17 species analysed, but some LGT might remain undetected due to our stringent phylogenetic filtering, and because we are only considering transfers among the predefined five grass clades.

Among the 17 species screened, LGT is observed in all functional groups (Figure 3). We detected LGT in wild species, but also in major crops. For instance, maize (*Zea mays*) has 11 LGT received from Chloridoideae and Paniceae, while wheat (*Triticum aestivum*) has 10 LGT received from Andropogoneae, Chloridoideae and Paniceae (Table 2). The LGTs may be beneficial for the crops, with transferred loci including some with functions related to abiotic stress tolerance and disease resistance (Table S2). Across all plant properties, LGT seems to be more abundant in perennial, rhizomatous and C4 species (Figure 3). A phylogenetic generalized least squares (PGLS) analysis was conducted to test for an effect of all traits while accounting for phylogenetic effects. For this we constructed a model to explain the absolute number of LGTs using nine traits as predictor variables (Table 1) and a time-calibrated phylogenetic tree retrieved from Christin *et al*., (2014). Initially, models were constructed for each predictor variable, with the amount of LGT shown to increase with the presence of rhizomes (p-value = 0.026, adjusted R^2^ = 0.243) and the number of genes tested (p-value = 0.038, adjusted R^2^ = 0.207). We subsequently performed a combined model with all explanatory variables to test for their joint effects. Iterative models were performed, removing the least significant variable until only significant variables remained (p-value <0.05). The PGLS analysis (combined adjusted R^2^ = 0.652) identified three characteristics that jointly increased the number of LGT: the number of genes tested (p-value < 0.001), the presence of rhizomes (p-value = 0.002), and the ploidy level (p-value = 0.006). Future studies should use larger sample sizes to definitely demonstrate the effects, but our analyses suggest that some categories of grasses are more likely to be involved in LGT.

**Fig. 3:**
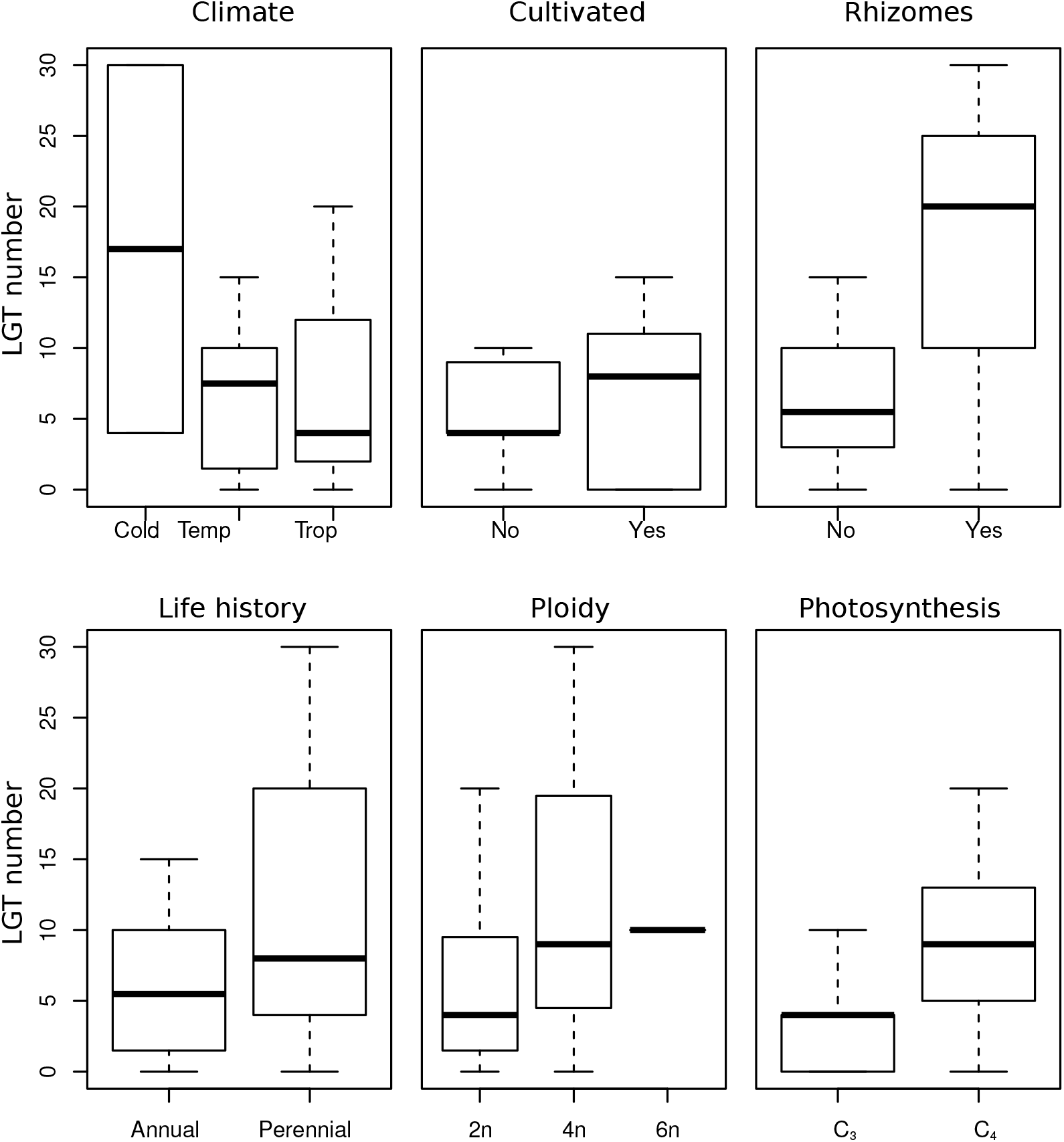
Numbers of lateral gene transfer (LGT) received by different categories of grasses. For each group, the distribution of LGT numbers is shown with box plots connecting the median and the interquartile range, with whiskers showing 1.5 x the interquartile range.

### 3.2 LGT are more commonly received from closely related species

Overall, some clades acted more frequently as donors (p-value < 0.01, Kruskal-Wallis test). Specifically, the Andropogoneae were the source of most transfers (Table 2). However, these were mainly received by members of Paniceae, which are the closest relatives of Andropogoneae in our dataset, and are also represented by the most genomes (Table 1). While these patterns suggest that LGT occurs more frequently among close relatives, directly comparing the rates is difficult because the clades vary in their number of species, number of genomes available and age. However, for a given clade of recipients, it is possible to compare the frequency of different groups of donors. We therefore focused on the identity of donors of LGT to Paniceae, the group with the highest number of complete genomes.

Seven Paniceae genomes were used in this study, and this increased sample size further allows to detect intra-Paniceae LGT. We therefore reported the number of LGT transferred from the Panicinae and Cenchrinae subgroups of Paniceae (each represented by two genomes) to other Paniceae, in addition to those received from other groups. In total, we identify 129 LGT across the seven Paniceae genomes, 35 of which were transferred from the Cenchrinae and Panicinae subgroups (Table 3; full results Table S4). When focusing on Paniceae recipients, some groups are more often LGT donors than others even after correcting for the number of species in each donor clade (p < 0.01, Kruskal-Wallis test). The number of LGT given per species decreases with the phylogenetic distance to Paniceae, reaching lowest levels in the BOP clade (Pooideae and Oryzoideae; Figure 4).

**Fig. 4:**
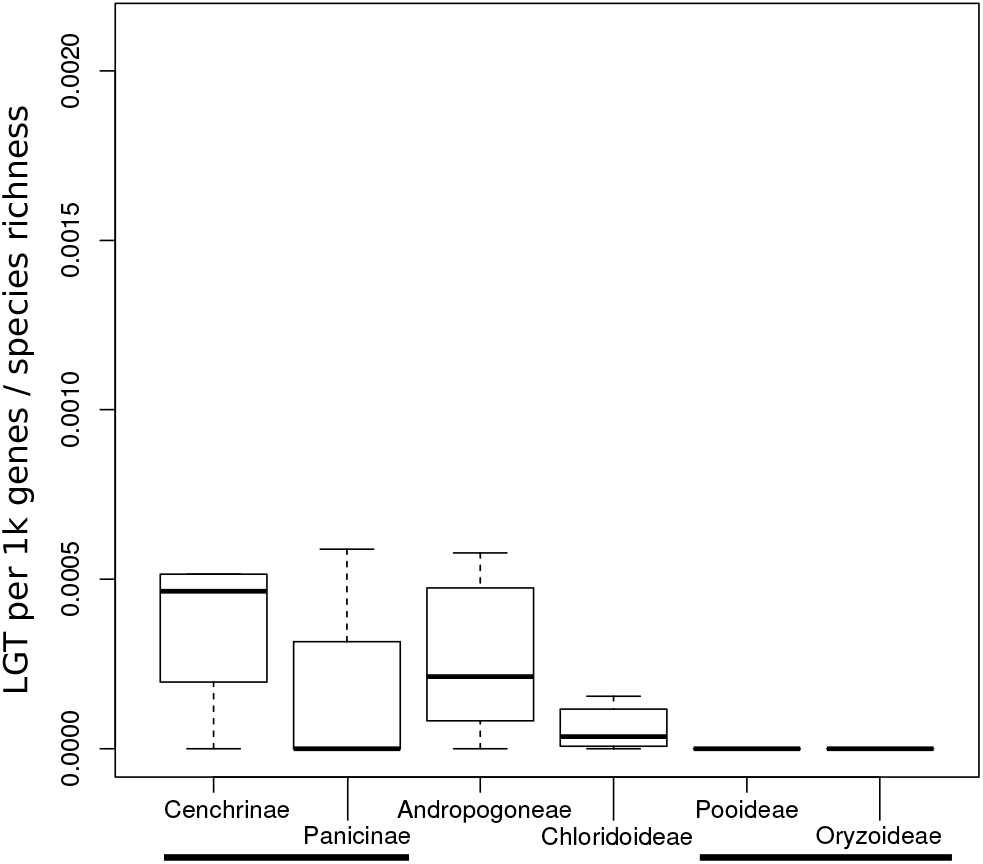
Number of lateral gene transfer (LGT) received by Paniceae species from different clades. The number of LGT in each Paniceae genome is corrected by the number of genes tested as well as the number of species in the group of donors. The phylogenetic distance increases from left to right, with equidistant clades joined by solid bars. Box plots show median, interquartile range and 1.5 x interquartile range.

**Table 3:** Number of lateral gene transfers (LGT) detected in Paniceae.

| Subgroup | Species | #<br>LGT | Pooid.<br>(3,698 sp.) | Ory.<br>(115 sp.) | Chlor.<br>(1,602 sp.) | Andro.<br>(1,202 sp.) | Cench.<br>(287 sp.) | Pani.<br>(157 sp.) |
| --- | --- | --- | --- | --- | --- | --- | --- | --- |
| Cenchrinae | *Cenchrus americanus* | 16 | 0 | 0 | 5 | 10 | - | 1 |
| Cenchrinae | *Setaria italica* | 7 | 0 | 0 | 0 | 7 | - | 0 |
| Panicinae | *Panicum hallii* | 8 | 0 | 0 | 6 | 2 | 0 | - |
| Panicinae | *Panicum virgatum* | 36 | 0 | 0 | 1 | 29 | 6 | - |
| Other | *Alloteropsis semialata* | 33(34) | 0 | 0 | 4 | 16 | 13(14) | 0 |
| Other | *Dichanthelium*<br>*oligosanthes* | 5 | 4 | 0 | 0 | 0 | 1 | 0 |
| Other | *Echinochloa crus-galli* | 23 | 0 | 0 | 3 | 7 | 8 | 5 |

The number of species in each clade is indicated in parentheses, with values from Soreng et al., (2015); Pooid. = Pooideae; Ory. = Oryzoideae; Chlor. = Chloridoideae; Andro. = Andropogoneae; Cench. = Cenchrinae; Pani. = Panicinae.

### 3.3 Ruling out alternative hypotheses

There are four main alternative hypothesis to LGT: [1] incomplete lineage sorting, [2] unrecognised parology, [3] hybridisation, [4] contamination, and [5] phylogenetic bias, such as convergent evolution. Below we present evidence reducing the likelihood of these alternative explanations.

[1] Incomplete lineage sorting: for a majority of the LGTs we detect (79.4%), the recipient genome also contains a native copy, which argues against incomplete lineage sorting as an alternative hypothesis. However, as pseudogenization of the native copy has been observed in cases where the LGT acts as a functional replacement (Dunning *et al*., 2019), their continued coexistence should not always be expected. The coexistence of native and laterally acquired orthologs permits us to compare patterns of synteny in multiple species to rule out unrecognised parology problems.
[2] Unrecognised parology: we used seven diploid species for this analysis, with at least one representative from each of the five groups. For each LGT detected in these seven species, we determined whether the genes from the other six species identified as orthologous in the phylogenetic tree were syntenic to the LGT or the native gene. In no instance was a gene from another model species syntenic with the LGT, with 76.3% being syntenic with the native copy, and 23.7% being syntenic to neither (Table S3). The synteny analyses therefore confirm that our phylogenetic trees identify true orthologs in most cases, and the phylogenetic patterns suggesting LGT cannot be explained by widespread unrecognised paralogy.
[3] Hybridisation: the patterns of synteny between the native and laterally acquired genes also argue against straightforward hybridisation through sexual reproduction and chromosomal recombination during the transfers, as already argued previously (Dunning *et al*., 2019). Indeed the LGTs appear to be inserted into the genome in random locations, often on different chromosomes as the native orthologs.
[4] Contamination: we rule out contamination as the source of the foreign DNA in the 13 genomes with LGT by confirming the presence of the laterally acquired DNA in multiple independent sequencing runs (Table S5 and Figure S1). For six of the reference genomes, ‘gold-standard’ datasets exists, i.e. whole-genome resequencing data sets for the same cultivar as the reference genome, but that were produced independently from the initial assembly project. For the remaining 7 species, only the whole-genome data used to generate the reference assembly exists, with all but one (*Leersia perrieri*) having multiple sequencing libraries/runs. For each dataset, we compare the genome-wide mean coverage for each gene to that of the identified LGT (Table S5 and Figure S1). All LGTs had sequencing data in the multiple datasets apart from one gene in *Z. mays*. For this species, we used seven datasets from the same cultivar that were produced independently in seven different labs. Only five out of these seven datasets supported the presence of the LGT *Zm00001d039537*, with the most parsimonious explanation being LGT variation between individuals, as previously documented in *Alloteropsis semialata* (Dunning *et al*., 2019). A majority of LGTs had coverage depths greater than the 5^th^ (97.0% of LGTs) and 2.5^th^ (99.0% of LGTs) percentile of coverage depth for all genes in the genome (Table S5 and Figure S1). Overall, these results confirm that contamination in the original reference genomes is not responsible for the presence of the LGT in the sequence datasets.
[5] Phylogenetic bias: convergent evolution or other systematic biases in the data could lead to gene/species tree discordance (Chang & Campbell, 2000). In addition to confirming the patterns with phylogenetic trees built on third positions of codons, we assessed the similarity between the recipient and donor species in non-coding DNA. The mapping of short-read data to four genomes confirmed a high similarity between the putative donor and recipient on intron sequences of LGT in addition to exons (Figure 2). It was however not possible to delimit with high precision the laterally acquired fragments detected here (as done for *A. semialata* in Olofsson *et al*., 2019), either because the transfers are too ancient or because we lack whole genome data for very close relatives of the donors. The observation of some intergenic regions with high similarity (Figure S2), together with intronic similarities (Figure 2), still rules out convergent evolution or other phylogenetic biases (e.g. long branch attraction) as being responsible for all detected cases of gene/species tree discordance.

## 4. Discussion

Lateral gene transfer is a potent evolutionary force capable of having a profound impact on the evolutionary trajectory of a species (Li et al., 2014; Cheng et al., 2019; Phansopa et al. In press). Here, we use grasses as a model system to investigate the factors that dictate the prevalence of LGT among plants. Using a combination of stringent phylogenetic and genomic analyses, we have identified a grand total of 170 genes (approximately 3.72 LGT per 10,000 genes; 135 in the first round of analyses and 35 among groups of Paniceae) that have been laterally transferred to 13 of the 17 complete grass genomes that were screened (Table 1 & Table 3). Our approach was developed to drastically reduce the amount of false positives, and is purposely very conservative. This enables us to minimise the effects of other evolutionary processes such as hybridisation and incomplete lineage sorting. As a result, the number of LGT identified is likely only a subset of those existing in the complete grass genomes. In addition, the phylogenetic filtering prevents us from detecting LGT from clades of grasses for which no genome is available. With the current sampling, at least 30% of the grass diversity is never considered as potential LGT donors (Soreng *et al*., 2015).

Our phylogenetic pipeline prevents us from detecting LGT happening among members of the same group of grasses, such as the numerous exchanges among lineages of Paniceae previously detected (Dunning *et al*., 2019). This is perfectly exemplified by the case of *A. semialata*, in which 59 LGT were previously detected (Dunning *et al*., 2019). Here, only 20 were identified when considering solely LGT among the five higher groups (Table 2), while a further 13 were detected when considering subgroups of Paniceae as potential donors (Table 3). The other LGTs previously identified for *A. semialata* were not detected in the present study because of slight methodological differences (e.g. less donor clades considered and sampling more genomes means greater multiple testing correction), and because we did not screen the flanking regions of LGTs for additional genes with high sequence similarity in the present study due to the larger scale of the present analyses, varying genome assembly contiguity, and a lack of whole-genome data for some donor species. Conversely, improvements in the initial screening approach (i.e. removing an initial read-mapping step from Dunning *et. al*., 2019 method) resulted in the identification of four novel LGT. These examples prove that our ability to detect LGT depends on many factors and strongly suggest that the LGTs we report here concern only a small fraction of those existing in grass genomes. Finally, our approach precludes the detection of older LGT that are shared by multiple individuals within a clade. Despite these limitations, we show that LGT is common in grasses, certain groups exchange more genes than others, the frequency of LGT appears to increase in rhizomatous species, and there may be a role of phylogenetic distance underpinning the LGT dynamics.

### 4.1 LGT occurs in all functional groups, and is especially prevalent in rhizomatous species

LGT is common in grasses and is observed in each of the five groups investigated here (Figure 1). We detected LGT in domesticated and wild species alike (Figure 3), although it is currently unknown whether the LGTs occurred before or after domestication and whether these genes are associated with agronomic traits. The genetic exchanges are not restricted to any functional category of grasses (Figure 3), and the ubiquity of the phenomenon provides some support for a breakdown in reproductive behaviour and illegitimate pollination as the mechanism responsible for the transfers as wind pollination is universal in this group. However, there is a statistical increase of the number of LGT in rhizomatous species and two of the three species with the highest numbers of LGT (*Alloteropsis semialata* and *Panicum virgatum*) are perennials that can propagate vegetatively via rhizomes (Table 1 & Table 3). These patterns suggest that root-to-rhizome contact (i.e. inosculation) provides an increased opportunity for retaining gene transfers, as the integration of foreign DNA in rhizome tissue means that any subsequent plant material regrown from these cells, including reproductive tissue, will contain the LGT. This hypothesis is compatible with previous reports of genetic exchanges following grafts (Stegemann & Bock, 2009). In this instance, LGT is similar to somatic mutations occurring in clonal species, as documented in the seagrass (*Zostera mariana*) where they can ultimately enter the sexual cycle (Yu *et al*., 2020). The genetic bottleneck and selection characterising rhizomes would further increase the chance of LGT retention, especially if these provide a selective advantage (Yu *et al*., 2020). However, we did not detect LGT in the third rhizomatous species we sampled (*Zoysia japonica*; Table 1). Increased species sampling, particularly for rhizomatous species represented by only three genomes in this study, is now needed to confirm our conclusions and precisely quantify the impact of growth form on the amount of gene transfers and how it interacts with other factors.

### 4.2 It is easier to acquire genes from close relatives

Within grasses, there is an effect of the phylogenetic distance on the number of transfers observed, as shown by the Paniceae receiving more LGT from closer relatives (Figure 4). This pattern mirrors that observed in prokaryotes (Popa & Dagan, 2011; Skippington & Ragan, 2012; Soucy *et al*., 2015) and insects (Peccoud *et al*., 2017), where the frequency of transfers is higher between closely related species. In prokaryotes, this effect is thought to result from more similar DNA sequence promoting homologous replacement of the native copy (Skippington & Ragan, 2012). This is unlikely to play a role in grasses as the LGTs are inserted in non-syntenic positions in the genome where they coexist with the native copy, often on different chromosomes (Table S3). However, stretches of DNA similar between the donor and recipient (e.g. transposable elements) may still be involved in the incorporation of the LGT onto the chromosomes, a hypothesis that can be tested when genome assemblies for donor species are available (e.g. *Themeda triandra;* Dunning *et al*. 2019). Alternatively, the effect of the phylogenetic distance might stem from the regulation of the LGT post acquisition, with genes transferred from closely related species more likely to share regulatory mechanisms. In such a scenario, the phylogenetic effect would reflect the utility of the LGT for the recipient species and therefore selection after the transfer rather than the rate of transfer. Overall, our analyses indicate that it is easier to either obtain LGT from close relatives or to use it after the transfers, thereby increasing the chance of selectively retaining it.

### 4.3 The role of overlapping distributions

In addition to genomic compatibility, the probability of co-occurrence in the wild might decrease with divergence time, explaining the observed effect of phylogenetic distance (Figure 4). Paniceae grasses generally grow in biodiverse savannas that are dominated by Andropogoneae species in wetter areas, and Chloridoideae grasses in drier habitats (Lehmann *et al*., 2019). The effects of phylogenetic distance are therefore confounded with those of biogeography, but specific examples indicate that biogeography can take precedence.

We observe some transfers between Pooideae and Paniceae, two groups that diverged >50 Ma, representing one of the earliest splits within this family (GWPGII, 2012). This indicates that LGT is possible across the whole grass family. In our dataset, the only recipient of these transfers is *Dichanthelium oligosanthes* (Table 2), a frost-tolerant grass from North America that inhabits colder areas than other members of the Paniceae (Studer *at al*., 2016). In cold regions, *D. oligosanthes* co-occurs with members of the Pooideae, and this biogeographic pattern likely facilitated exchanges between the two groups of grasses. However, given the difficulties of identifying the donor to the species level (or even genus) with the current data, we can not be sure that the specific donor and *D. oligosanthes* co-occur. As more whole-genome datasets become available for the diverse Pooideae, co-occurrence between the donor and recipient species can be directly tested.

Biogeography might also be responsible for differences in the identity of the LGT donors between the two closely related *Panicum* species. Indeed, a majority (75%) of LGT in *Panicum hallii* were received from Chloridoideae, while a majority (81%) of those in *Panicum virgatum* were received from Andropogoneae (Table 3). This pattern mirrors the dominant grassland type (Chloridoideae vs. Andropogoneae) for a majority of the range of each of the two species, and the area from which the individual for the genome assembly was sampled (Lovell *et al*., 2018; Lehmann *et al*., 2019).

Quantifying the effects of biogeography as opposed to other factors requires identifying the donor to the species level and a detailed description of the spatial distribution of each grass species, including their abundances. Indeed, the likelihood of encounters will increase with the number of individuals of the donor species and not just its presence. Such ecological datasets coupled with genomic data for a large numbers of grasses will be key to future studies of LGT dynamics in this group of plants.

### 4.4 Conclusions

Using stringent phylogenomic filtering, we have shown that lateral gene transfer (LGT) is a widespread process in grasses, where it occurs in wild species as well as in widely cultivated crops (e.g. maize and wheat). LGT does not appear restricted to particular functional types, although it seems to increase in rhizomatous species, where vegetative growth offers extra opportunities for gene transfers into the germline. In addition, we show that the amount of successful transfers decreases with phylogenetic distance. This effect of the phylogenetic distance might result from increased genomic compatibility among more related groups. Alternatively, groups that diverged more recently might be more likely to co-occur, offering more opportunities for genetic exchanges. Indeed, biogeography seems to have an overall effect on the frequency of LGT, as the only species that received genes from a group that diverged more than 50 million years ago is the one with an overlapping distribution following a relatively recent niche shift. Overall, our study shows that LGT occurs in a variety of grasses, and the frequent movements of functional genes can strongly impact the evolution of this group of plants.

## 5. Acknowledgements

This work was funded by a European Research Council Grant (ERC-2014-STG-638333) and a Royal Society Research Grant (RGF\EA\180247). P.-A.C. is supported by a Royal Society University Research Fellowship (URF\R\180022). L.T.D is supported by a Natural Environment Research Council Independent Research Fellowship (NE/T011025/1).

## 6. Authors’ contributions

All authors designed the project. LTD, SGSH and PR conducted the analyses. LTD, PAC and SGSH wrote the paper, with the help of PR.

