## Supplemental Figures, Tables & Datasets for "Widespread lateral gene transfer among grasses": Supporting Information.pdf

**Dataset S1:** Nucleotide alignments.

**Dataset S2:** LGT maximum-likelihood trees, 85 taxa.

**Dataset S3:** LGT maximum-likelihood trees, 85 taxa 3rd codon position.

**Dataset S4:** LGT maximum-likelihood trees, including short-read data.

**Figure S1:** Coverage plots from contamination analysis.

**Figure S2:** Laterally acquired fragment in *Setaria italica* genome.

**Table S1:** Data sets used.

**Table S2:** Detailed summary of results for the main analysis**.**

**Table S3:** Results of the synteny analysis.

**Table S4:** Detailed summary of results for the within Paniceae analysis**.**

**Table S5:** Coverage data used for the contamination analysis.

**Figure S1:** Determining if contamination was the source of the lateral gene transfers (LGT) identified in the grass reference genomes. Each graph represents a different NCBI Sequence Read Archive data set with the species and the accession numbers indicated. The black histogram represents the mean coverage for all genes in the genome. The sold blue line represents the 2.5th mean coverage percentile, and the dashed blue line the $5^{th}$ percentile. The red lines indicate the mean coverage of the indivdual LGTs detected in the genome. The figure consist of four panels.

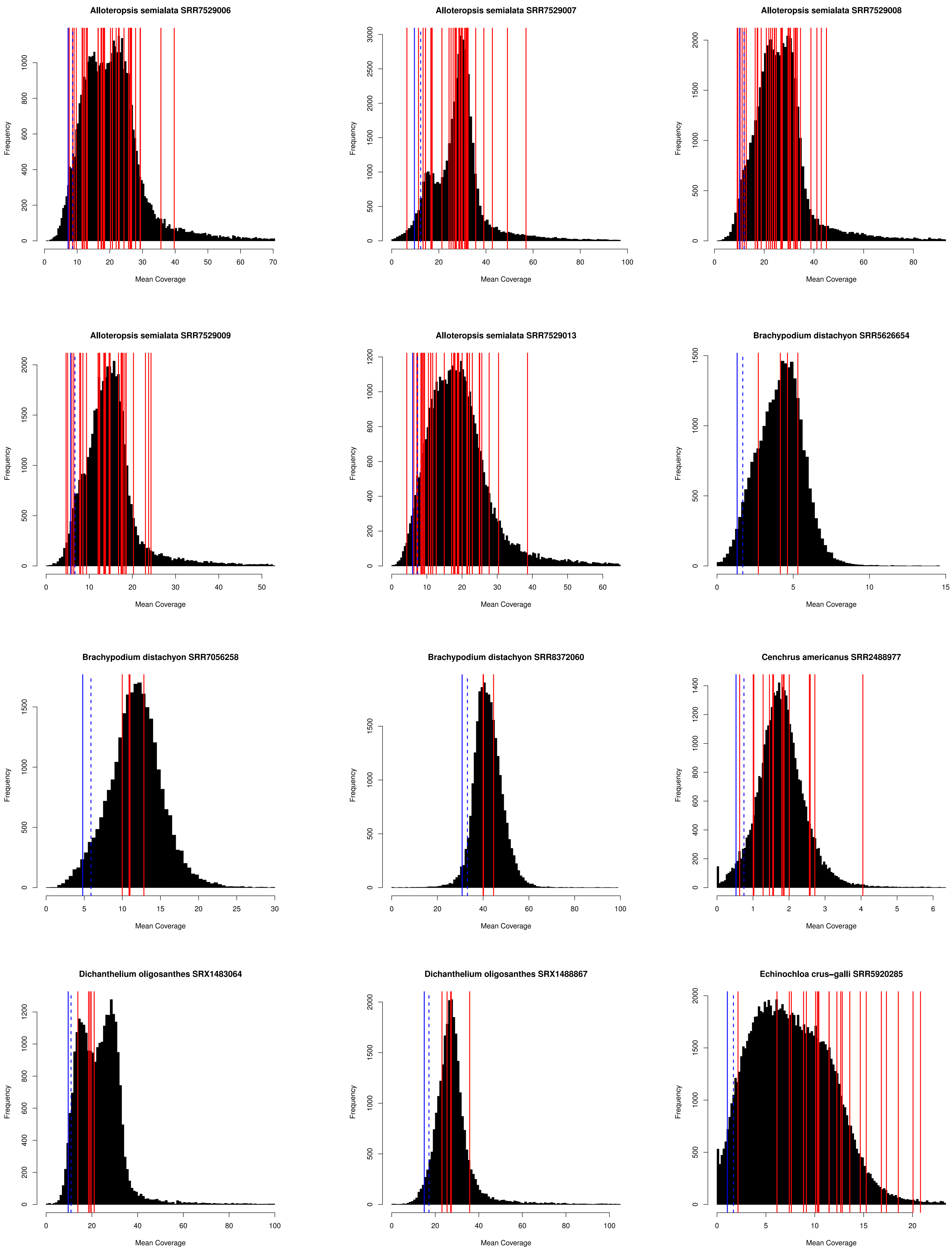
Alloteropsis semialata SRR7529006
Alloteropsis semialata SRR7529007
Alloteropsis semialata SRR7529008
Alloteropsis semialata SRR7529009
Alloteropsis semialata SRR7529013
Brachypodium distachyon SRR5626654
Brachypodium distachyon SRR7056258
Brachypodium distachyon SRR8372060
Cenchrus americanus SRR2488977
Dichanthelium oligosanthes SRX1483064
Dichanthelium oligosanthes SRX1488867
Echinochloa crus-galli SRR5920285
Frequency
Mean Coverage

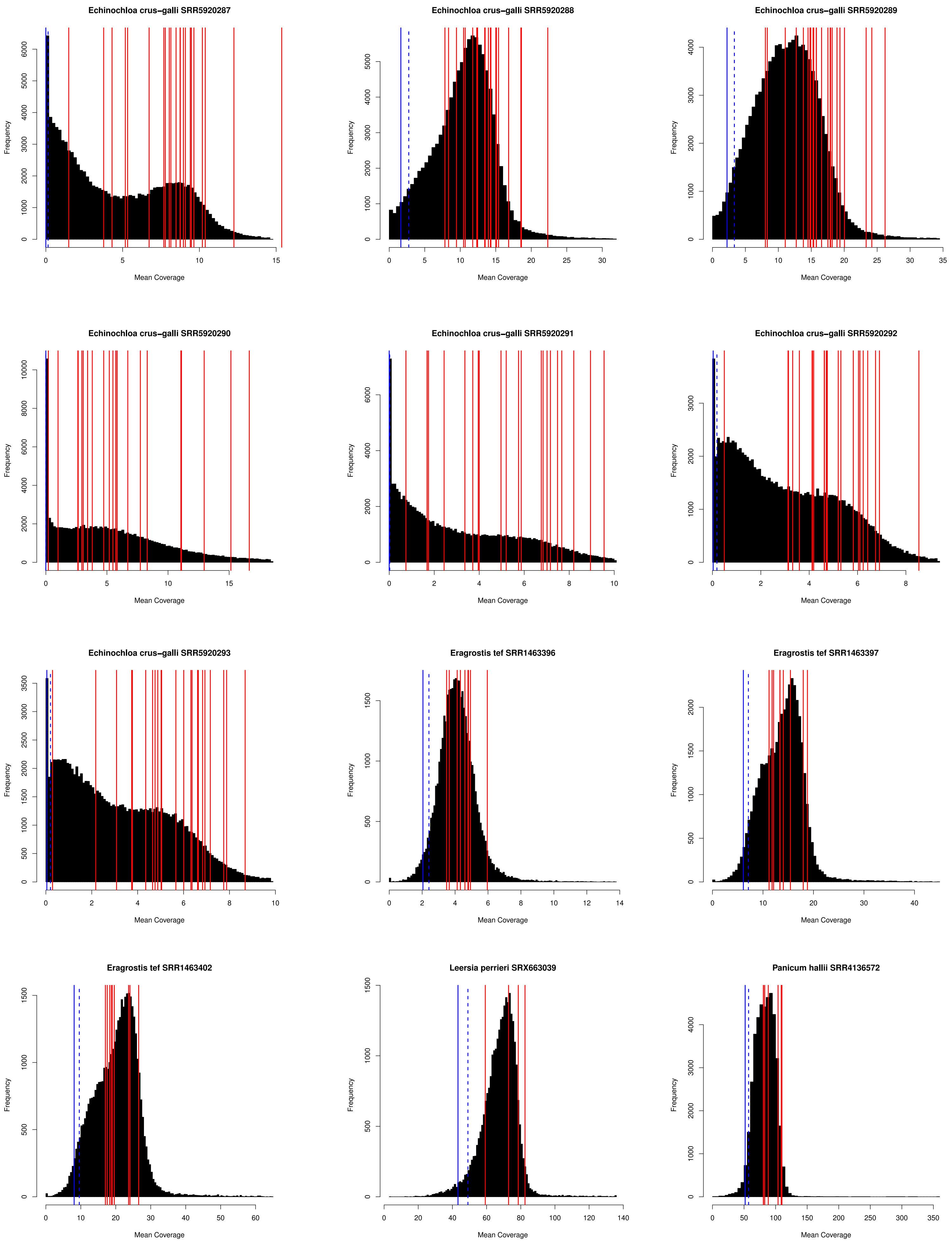
Echinochloa crus−galli SRR5920287
Echinochloa crus−galli SRR5920288
Echinochloa crus−galli SRR5920289
Echinochloa crus−galli SRR5920290
Echinochloa crus−galli SRR5920291
Echinochloa crus−galli SRR5920292
Echinochloa crus−galli SRR5920293
Eragrostis tef SRR1463396
Eragrostis tef SRR1463397
Eragrostis tef SRR1463402
Leersia perrieri SRX663039
Panicum hallii SRR4136572
Frequency
Mean Coverage

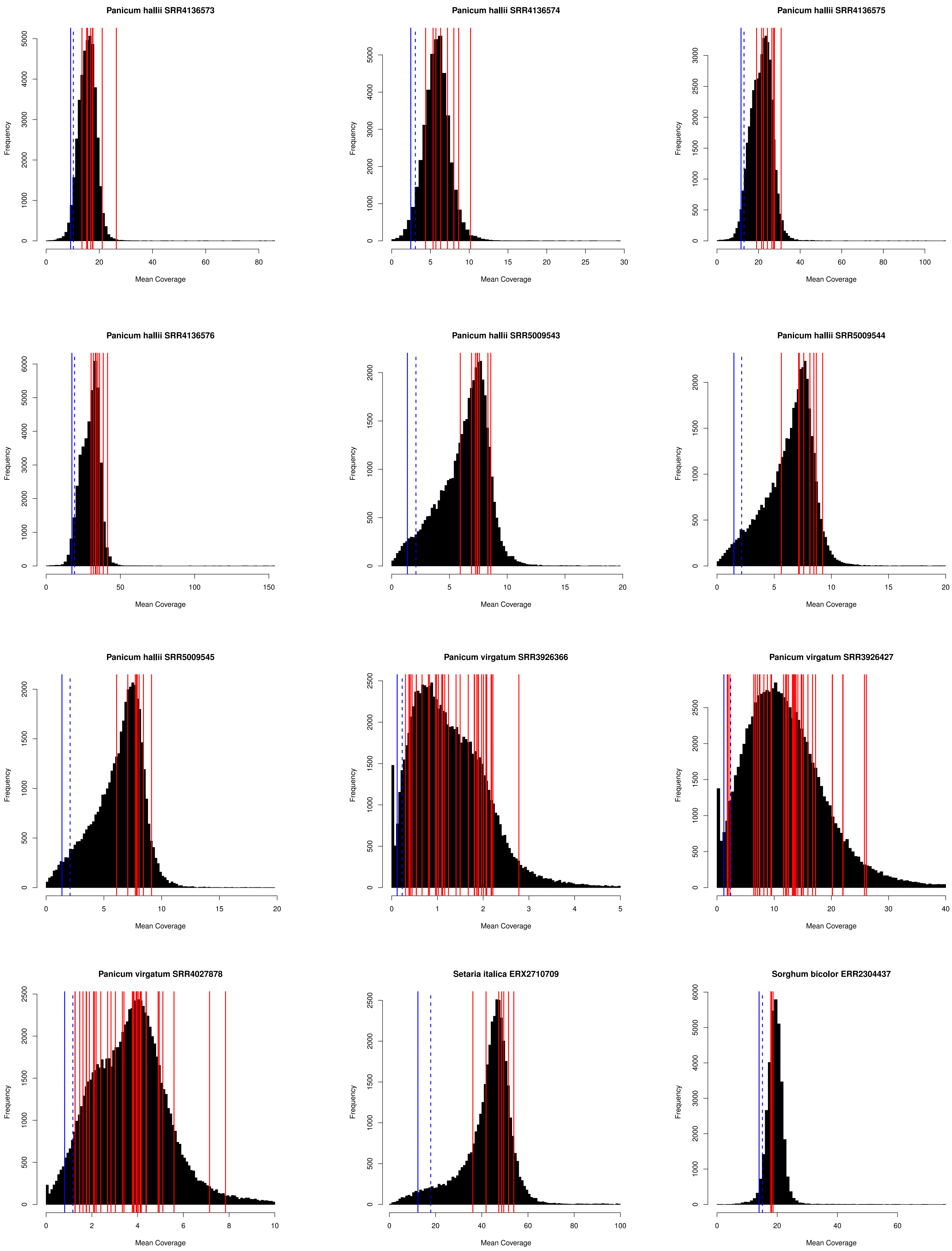
Panicum hallii SRR4136573
Panicum hallii SRR4136574
Panicum hallii SRR4136575
Panicum hallii SRR4136576
Panicum hallii SRR5009543
Panicum hallii SRR5009544
Panicum hallii SRR5009545
Panicum virgatum SRR3926366
Panicum virgatum SRR3926427
Panicum virgatum SRR4027878
Setaria italica ERX2710709
Sorghum bicolor ERR2304437
Frequency
Mean Coverage

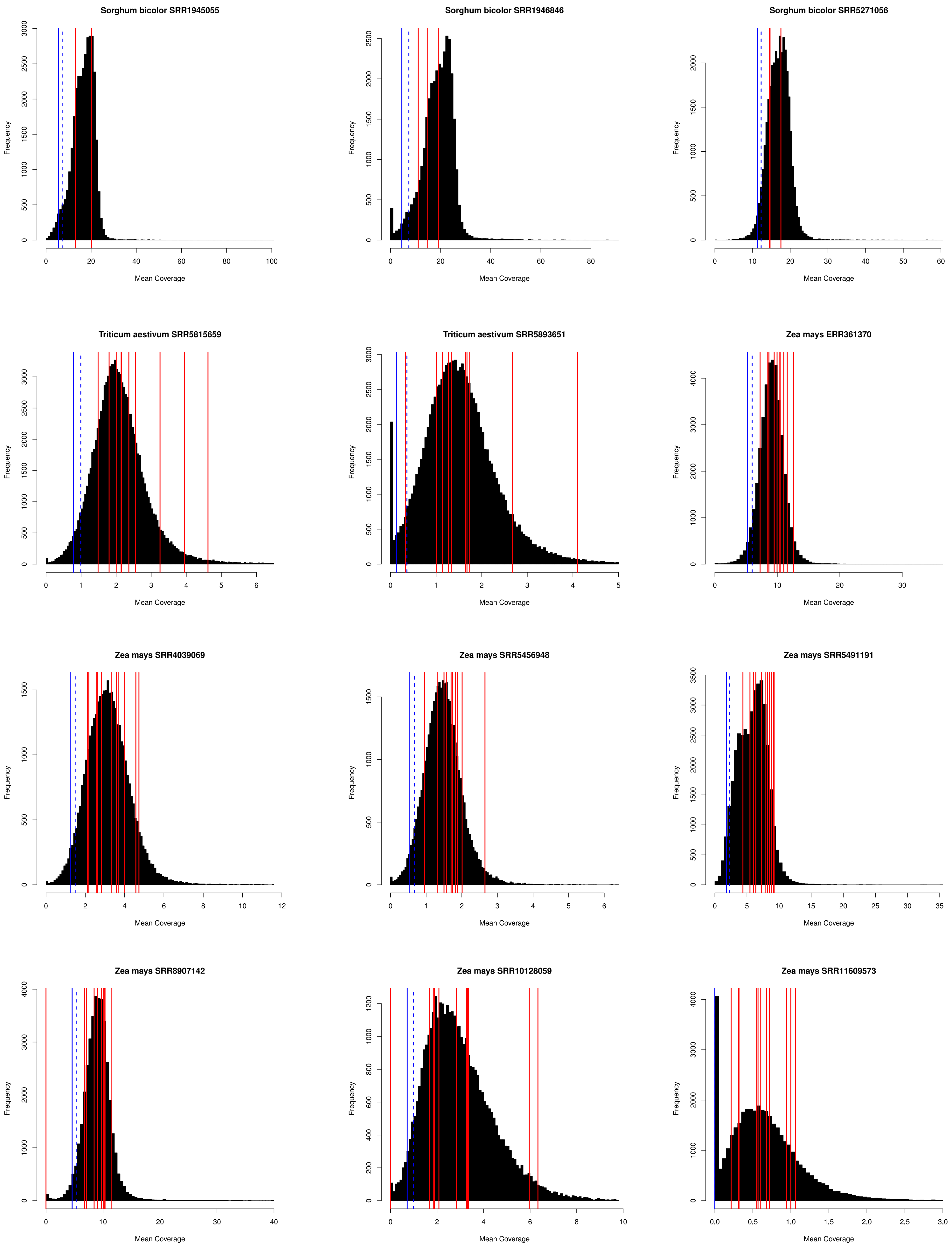
Sorghum bicolor SRR1945055
Sorghum bicolor SRR1946846
Sorghum bicolor SRR5271056
Triticum aestivum SRR5815659
Triticum aestivum SRR5893651
Zea mays ERR361370
Zea mays SRR4039069
Zea mays SRR5456948
Zea mays SRR5491191
Zea mays SRR8907142
Zea mays SRR10128059
Zea mays SRR11609573
Frequency
Mean Coverage

**Figure S2:** Three laterally acquired genes (position shown in green) acquired from an Andropogoneae species are found in a row on chromosome III of the *Setaria Italica* genome. The results of mapping high-coverage data for the Andropogoneae species *Sorghum bicolor* to this region is shown, with valid read alignments having a nucleotide identity ≥ 90%. The *S. bicolor* data maps to both genic and intergenic regions. Two of these loci (Si024038m & Si024806m) are recent duplicates, hence why a majority of the mapping data is low-quality in these regions.

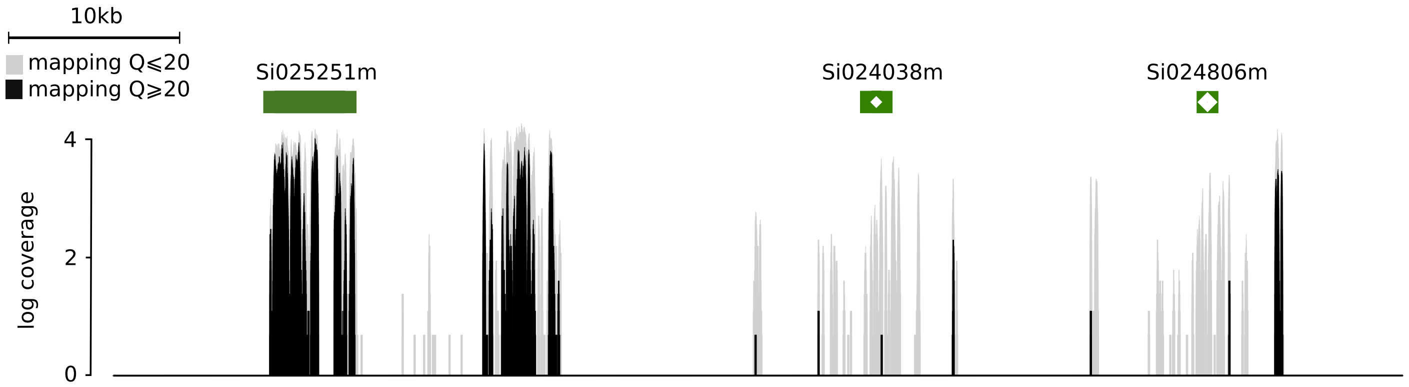
